# Late-life mortality plateaus through error, not evolution

**DOI:** 10.1101/410134

**Authors:** Saul Justin Newman

## Abstract

This study highlights how the mortality plateau in Barbi *et al.* [1] can be generated by low frequency, randomly distributed age misreporting errors. Furthermore, sensitivity of the late-life mortality plateau in Barbi *et al.* [1] to the particular age range selected for regression is illustrated. Collectively, the simulation of age misreporting errors in late-life human mortality data and a less specific model choice than that of Barbi *et al*. [1] highlight a clear alternative hypothesis to the cessation of ageing.

## Text

Barbi *et al.* [1] proposed that late-life plateaus were caused by the cessation of ageing. However, Barbi *et al.* correctly highlighted an alternative hypothesis: the potential for age misreporting and cohort blending errors to generate late-life plateaus. Despite raising this hypothesis, the effect of low-frequency age misreporting errors was not actively addressed.

Therefore, late-life plateaus observed in the 1904 cohort were compared to the effect of simulated, low frequency age-reporting errors introduced into log-linear models of mortality fit to age range of 65-80 used in Barbi *et al.* [1].

Random, normally distributed age-coding errors were seeded into these synthetic cohorts at age 50, with a probability ranging from p = 10^−3^to p = 10^−6^(Fig 1a). Age misreporting errors generated late-life plateaus at frequencies below 1 in 500 in the model presented by Barbi *et al.* (Fig 1a-b). Furthermore, hazard rates resulting from these random errors closely resembled the late-life plateau of Barbi *et al*. (Fig 1a) and required no biological or evolutionary explanation.

**Figure 1.**
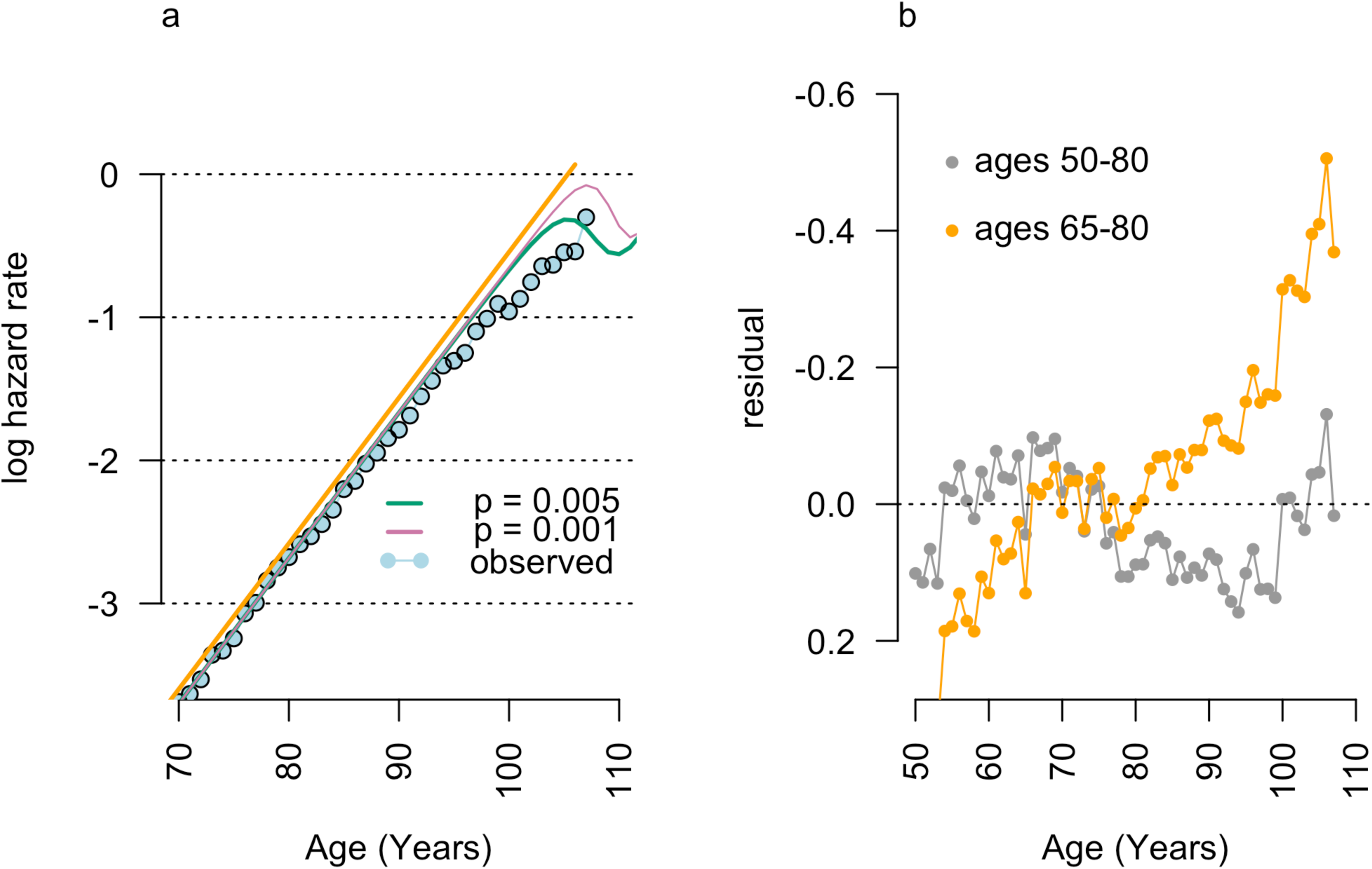
Generation of late-life mortality plateau by random errors. The introduction of symmetrically distributed age-coding errors **a,** into a log-linear model (orange line) fit to the 1904 cohort data used in Barbi *et al.* [1], generates late-life mortality patterns (green and pink lines) similar to observed data (blue points). Residuals from observed data **b** illustrate the skewed residuals in the model presented by Barbi *et al.* [1], compared with more representative regressions of hazard rates such as ages 50-80 shown (grey).

The apparent size of the late-life plateau in Barbi *et al.* [1] is characterized in comparison to a ‘best-estimate trajectory’ model, fit to data from a specific age range. The poor fit of this mid-life model to late-life data is used as justification for fitting separate late-life models. However, the choice of age range used to fit log-linear models has a large effect on the size and existence of late-life mortality plateaus (Fig 2).

**Figure 2.**
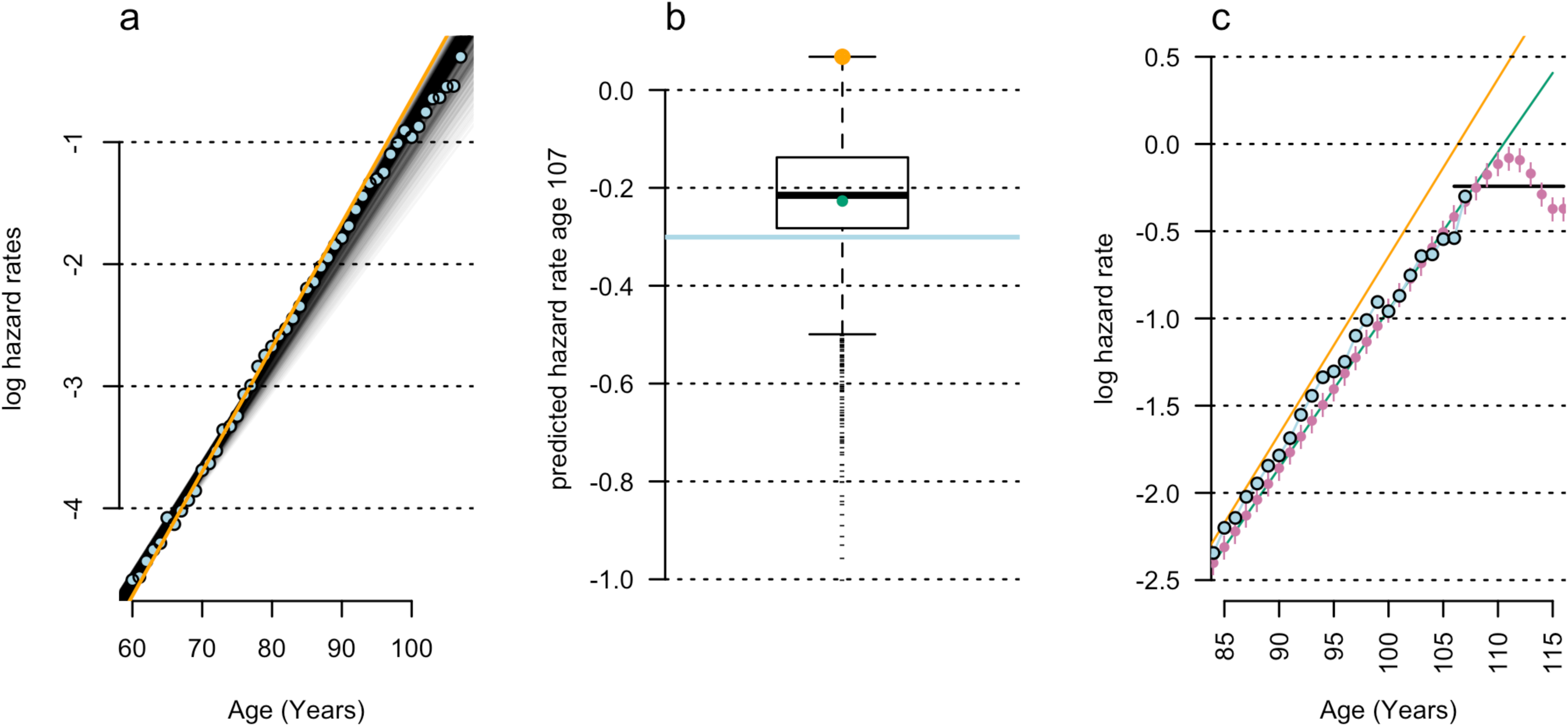
Effect of model selection on the size of apparent mortality plateaus. a. Observed hazard rate data (blue) from Barbi *et al.* [1], fit by log-linear hazard rate regressions for 861 diverse age ranges (grey lines). The Barbi *et al.* [1] age range (orange) produces the largest late-life mortality plateau and **b** produces the greatest over-estimate of observed data (orange cross) at advanced ages (blue line, age 105 shown). Seeding random errors into other representative models, *e.g.* the 50-80 year regression (green point in **b,** green line in **c**), produces **c** late-life mortality deceleration (pink points; p = 2e^−4^) and constant hazard rate regressions (black line) past age 105.

Barbi *et al.* selected an age range of 65-80 years [1]. This modeling choice was compared to log-linear models fit to mortality data using the 1904 Italian cohort data [2] used by Barbi *et al*. [1], fit to all possible age range combinations starting between ages 45-65 inclusive and ending between ages 70-107 inclusive. Of the 861 age-range combinations tested, the model selected by Barbi *et al.* generated the single largest late-life mortality plateau (Fig 2b-c). This model choice also provided the worst fit of mid-life (age 50) and late-life mortality patterns (Fig 2b).

Re-fitting these models to other age ranges reduced the apparent deviation of late-life data from mid-life patterns of mortality (Fig 1), and reduced or eliminated the late-life plateau in these data. Furthermore, models fit to alternative age ranges have a greatly reduced threshold for age-misreporting errors to cause late-life mortality deceleration and plateaus. For example, artificial mortality plateaus are produced by age misreporting rates below 1 in 10,000 if a log-linear model fit to ages 50-80 is used (Fig 2c). If these error patterns are fit by log-linear models using the same parameters as Barbi *et al.* [1], constant-hazard mortality ‘plateaus’ are produced (Fig 2c).

Simulated error rates were low compared to observed rates of manual entry errors. For example, double-entry of clinical trial data has error rates of p = 10^−3^ [3]. The pattern of late-life mortality deceleration shown in Fig 1a (pink line) results from random errors generated at half this rate. However, birth certificate data in the Barbi *et al.* data constitute hand-written records generated by a cohort with 32% literacy rates [4] and 9.6 months of education on average [5]. For error-generated late-life plateaus to be excluded from the data in Barbi *et al*., this cohort would have to achieve error rates 2-to 1000-fold lower than that observed in clinical trials. This seems unlikely.

Finally, claims by Barbi *et al*. that age data are validated by documents, and therefore ‘real’, should be viewed in the context of previously validated longevity claims. For example, Carrie White successfully claimed to be the world’s oldest and then second-oldest person. Documents certifying her age endured global scrutiny for 24 years. Her record was verified and accepted by the Gerontology Research Group and the Guinness world book.

The Carrie White record was retracted in 2012 after it was shown to be the result of a clerical error that inflated her apparent age from 102 to 116. A single age-coding error by a mental institution worker was copied to all later documents, and validated. Absent the survival of this otherwise obscure document, the Carrie White record would stand as a document-validated supercentenarian.

All supercentenarian data in Barbi *et al*. are susceptible to similar, but undetected, errors. In similar Gerontology Research Group data, 8% of all supercentenarian cases were found to be errors [6], and the potential for many other undocumented cases remains. As such, asserting that supercentenarian data are clean because they have passed document-based validation is unfounded.

The capacity for data entry and age inflation errors provides a sufficient model to explain late-life mortality patterns observed by Barbi *et al.* without requiring a cessation of ageing. This suggests the late-life mortality plateau observed was a result of errors, not biology.

